# Volatilized ammonia supports extremophilic cave ecosystems with unusual nitrogen isotopic signatures

**DOI:** 10.1101/2025.11.19.689070

**Authors:** Mackenzie B. Best, Scott D. Wankel, Heather V. Graham, Jennifer C. Stern, Jennifer Macalady, Maurizio Maniero, Stefano Recanatini, Sandro Mariani, Ilenia D’Angeli, Ilaria Vaccarelli, Daniel S. Jones

## Abstract

The sulfidic Frasassi cave system hosts a robust, subterranean ecosystem based on microbial lithoautotrophy. Curiously, acidic biofilms forming above degassing sulfidic cave streams, and the invertebrates that feed on them, are extremely depleted in nitrogen-15 (δ^15^N values less than -20‰). In this study, we tested the hypothesis that these low δ^15^N values result from the volatilization, trapping, and uptake of ammonia degassed from the circumneutral streams. We found that dissolved ammonium in the streams had δ^15^N values near +3‰, whereas NH_3_(*g*) in the cave atmosphere above streams exhibited δ^15^N values as low as -27‰, consistent with fractionation by NH_3_ volatilization. Extremely acidic condensation droplets on cave walls efficiently trapped airborne NH_3_, accumulating up to 4 mM NH_4_^+^ with δ^15^N values as low as -29‰, thereby confirming volatilized and trapped ammonia as the primary N source to cave wall biofilms and the extensive subsurface ecosystem they support. Airborne ammonia trapping represents a novel mechanism for biological N acquisition and provides abundant N for growth in an extreme subsurface environment that otherwise receives very limited nutrient input.

**Significance:** Nitrogen is one of the most abundant elements in organic molecules, and is a key limiting nutrient in many ecosystems. We showed that acidic microbial biofilms in sulfidic caves scavenge trace amounts of airborne ammonia, enabling microbial primary production and supporting associated food webs, including animals, in an environment where nitrogen is otherwise extremely scarce. This process represents a novel mechanism of biological nutrient acquisition and results in biomass and organic matter more depleted in the heavy isotope of nitrogen (^15^N) than almost all other biological materials on Earth. Extreme ^15^N isotope depletion is therefore a potential signature of acidic underground ecosystems on Earth and other planetary bodies.

## Introduction

As a major component of amino acids, nucleic acids, and other essential biomolecules, nitrogen (N) is fundamental for all known life forms, and its availability often limits biological productivity (1, 2). While abundant in Earth’s atmosphere, dinitrogen (N_2_) is inaccessible to all but specialized diazotrophic microorganisms, so organisms generally rely on other more bioavailable forms of N such as nitrate, ammonia, or organic N (3–5). Due to its importance in biological processes on Earth and reactivity in atmospheric and marine settings, nitrogen has been identified as an important target in the search for extraterrestrial life (6, 7). Earth environments with unusual N cycling and signatures are thus valuable for developing N-based biosignatures that could be used to evaluate the habitability of extraplanetary bodies (e.g., (8, 9). Earth’s shallow subsurface provides analog environments where microbial-dominated ecosystems adapt to severe nutrient limitation (10, 11) and sometimes depend on atypical nitrogen sources and cycling (12–14).

Sulfuric acid caves are hotspots for life in Earth’s subsurface. These caves contain ecosystems based on lithoautotrophy that form where deep-seated, hydrogen sulfide (H_2_S)-rich groundwaters are exposed to oxygen, resulting in a process called sulfuric acid speleogenesis (15). Sulfide-oxidizing microorganisms take advantage of the chemical energy at this mixing zone, catalyzing the aerobic oxidation of H_2_S and generating sulfuric acid (H_2_SO_4_) (Eq. 1) and elemental sulfur (S^0^) (Eq. 2), which can then be further oxidized to H_2_SO_4_ (Eq. 3). Below the water table, H_2_S oxidation is driven by microorganisms in streams and lakes that form distinctive white mats rich in S^0^ (16, 17). Above the water table, H_2_S_(*g*)_ that degasses from turbulent springs and streams into the cave atmosphere supports extremely acidic (pH < 2) sulfide-oxidizing communities on cave walls and ceilings (Fig. 1a) (18–20). Where H_2_SO_4_ aggressively dissolves the surrounding cave limestone, massive deposits of gypsum (CaSO_4_•2H_2_O) form on cave walls and ceilings as corrosion residues (Eq. 4) (21–23) (Fig. 1a).

**Figure 1.**
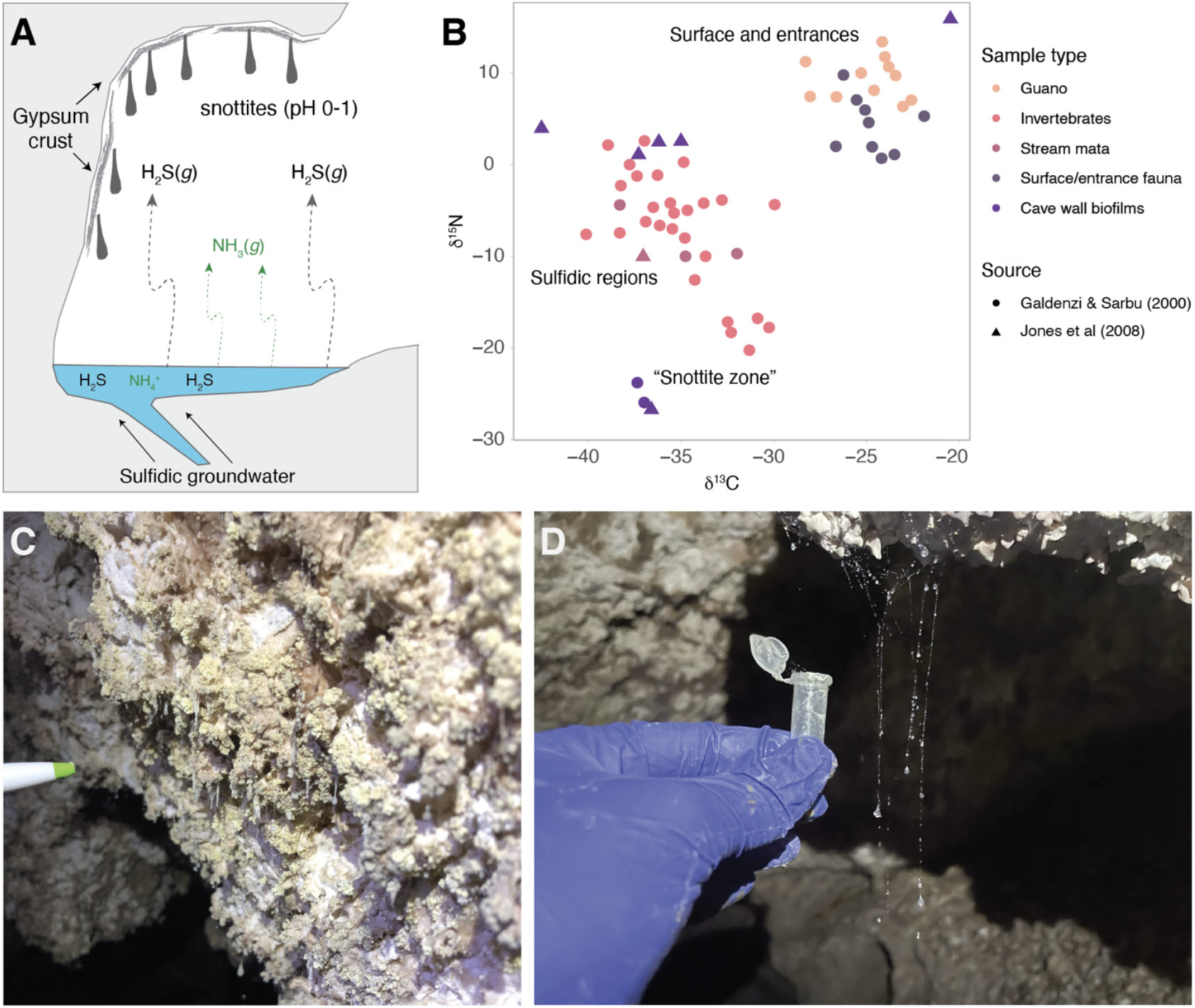
Panel A is a schematic depicting the processes occurring near degassing sulfidic cave streams, including the source of H_2_S and NH_3_ to the cave atmosphere. Panel B shows δ^15^N and δ^13^C values of organic material from the Frasassi Caves from previous work (27, 38). Panels C and D are photos of extremely acidic cave wall biofilms and spiderweb droplets that were sampled for this study.

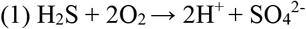

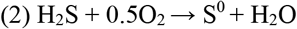

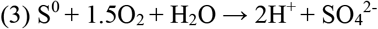

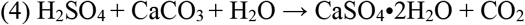

The lithoautotrophic nature of sulfidic cave ecosystems was first demonstrated in Movile Cave, Romania, where carbon and nitrogen isotope ratios (δ^13^C and δ^15^N, respectively) of organic matter at the sulfidic water table are isotopically distinct from surface-derived materials (24, 25). Subsequent work in the Frasassi Caves in central Italy revealed similar patterns, with low δ^13^C and δ^15^N values in organics collected near the sulfidic water table (26–28) and extremely acidic wall biofilms and associated invertebrates exhibiting unusually low N isotope ratios (Fig. 1b) (19). Very low δ^15^N values were also reported from Lower Kane Cave, Wyoming, USA, which were speculated to arise from NH_3(*g*)_ volatilization from the sulfidic aquifer and subsequent scavenging in sulfuric acid droplets (29). The accumulation of NH_3_ on cave walls is especially significant in the oligotrophic context of the cave habitat, because the acidic biofilms do not contain genes for N_2_ fixation (30, 31) and are not exposed to drip waters that could supply dissolved N compounds. Therefore, we measured the concentration and nitrogen isotopic composition of NH_3(*g*)_ and NH_4_^+^ in cave air, condensation droplets, and streams at Frasassi, in order to test the hypothesis of Stern et al. (2003) (29) that NH_3_ volatilization and scavenging provides a unique N supply mechanism that supports acidophile subsurface communities. Our results explain the extremely low δ^15^N values of subaerial sulfidic cave biofilms, mineral deposits, and invertebrates, and shed new light on N cycling in extremophilic subsurface environments.

### The Frasassi Caves

The Frasassi Caves are located in the northeastern Apennine Mountains in the Marche region of central Italy. Hosted in the core of a NNE-verging anticline in Jurassic limestone, the cave system has over 30 km of solutional passages developed in several levels.

The lowermost and youngest levels are actively forming at the sulfidic water table, where high concentrations of H2S in the air and water drive ongoing sulfuric acid speleogenesis (32). These passages are lined with acidic gypsum crusts on the walls and ceilings, and sulfidic streams and lakes occur where H_2_S-rich waters rise from a deep-seated aquifer. Cave streams are cold (13-14 °C), circumneutral (pH 6.9-7.4), and contain up to 600 µM dissolved sulfide and up to 175 µM dissolved ammonium (21, 33, 34). The sulfide is likely produced during microbial sulfate reduction in underlying Triassic evaporites (32). Higher cave levels represent progressively older passages that originally formed at the sulfidic water table but were abandoned and uplifted by tectonic activity (35, 36). These older levels show evidence of relict sulfuric acid corrosion in the form of massive gypsum “glaciers” that persist in places that are protected from dissolution by infiltrating meteoric water.

Previous research has shown that life near the sulfidic water table at Frasassi is supported by lithoautotrophic primary production (26–28), and includes endemic macroinvertebrates, some with lithoautotrophic symbionts (37). Very low δ^15^N isotope values (-24 to -26.5‰) have been reported from analyses of materials immediately adjacent to cave streams, such as biovermiculations and biofilms called snottites (19, 38) (Fig. 1b). Snottites—drip-shaped, hanging microbial biofilms abundant in areas of high H_2_S_(g)_ flux (Fig. 1c)—are dominated by sulfide-oxidizing microorganisms, which produce sulfuric acid and drive the exceptionally low pH (0 to 2) (30, 39). Biovermiculations, patterned wall formations resembling termite trails found on limestone surfaces both close to and far from the sulfidic water table (Supplementary Fig. S1), host more diverse communities and less acidic pH values (3 to 7) (38).

## Results

### Reduced nitrogen species in cave air and water

In addition to hydrogen sulfide, Frasassi cave streams also carry dissolved ammonium (NH_4_+) and ammonia (NH_3_) (Fig. 1). We sampled sulfidic stream water at three sites within the Frasassi cave system: Grotta Bella (GB), Pozzo dei Cristalli (PC), and Ramo Sulfureo (RS) (Fig. S2). Across two years, NH_x_ concentrations ranged from 80.7-158.0 µM NH_x_ (where NH_x_ refers to the combined NH_3_ + NH_4_^+^ pool), and pH ranged between 7.10 and 7.48 (Table S1). Under these conditions, the NH_4_^+^ species is far more abundant than the NH_3_ species (Eq. 5), and the expected concentrations of NH_3(g)_ in equilibrium with these waters (Eq. 5 and 6) range from 2 to 9 parts per billion by volume (ppbv) (Table S1).

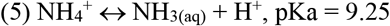

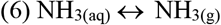

To capture and quantify NH_3(g)_ in the cave atmosphere, we deployed both passive and active samplers at varying distances from cave streams, ranging from 0.5 m, where H_2_S was readily detectable, up to >15 m from the cave streams where H_2_S_(g)_ was undetectable (Table S2). Total NH_3(g)_ concentrations in cave air ranged from 0.6 to 2.5 parts-per-billion by volume (ppbv), slightly lower than expected equilibrium values (Table S3). While the highest concentrations were in air samples collected closest to the sulfidic stream, no systematic variations in concentration with distance were observed (Table S3).

We also sampled NH_4_^+^ in condensation droplets (Fig. 1d), which ranged in size and appearance, with some clear and some cloudy white. Most droplets had pH values <1, except for several collected furthest from the sulfidic stream that were pH 4. Given the low pH of these droplets, NH_3(g)_ would readily protonate to NH_4_^+^ upon contact, thereby trapping nitrogen as NH_4_+. Acidic droplets collected within 1.5 m of cave streams contained an average of 2.23 mM NH_4_^+^ with concentrations reaching as high as 4.33 mM NH_4_^+^ (Fig. 2). In contrast, droplets collected 7.5-12.5 m from streams contained NH_4_^+^ ranging from 0.05 to 0.26 mM, and droplets collected 27.5 m away from streams contained an average of 0.018 mM NH_4_^+^ (Fig. 2, Table S4).

**Figure 2.**
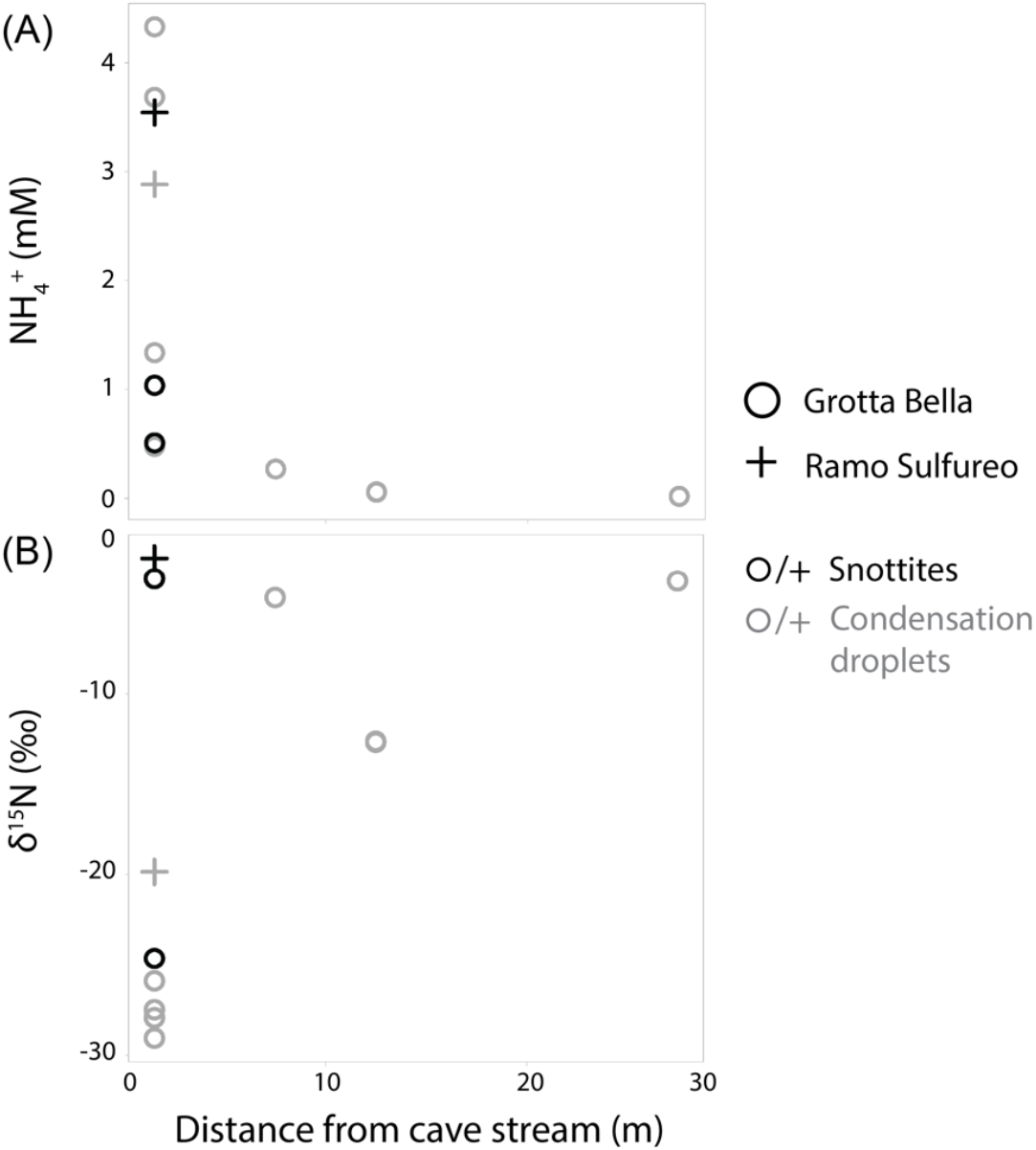
Ammonium (NH_4_^+^) concentrations (upper) and δ^15^N values (lower) in spiderweb and snottite droplets collected from sites GB and RS with distance from the degassing sulfidic cave stream. Shape corresponds to sample site, and color to sample type.

### Stable isotopic composition of cave ammonia and ammonium pools

In cave stream waters, average δ^15^N_NHx_ was +3.3‰, with values ranging from +0.3 to +4.7‰. This was consistent across sampling locations: δ^15^N_NHx_ in stream water ranged from +0.8 to +4.7‰ at site PC, from +0.3 to +4.4‰ at site RS, and was consistently +4.3‰ at site GB (Table S1).

We also measured the isotopic composition of airborne NH_3(*g*)_ collected using both passive and active samplers. Passive samplers, which used citric acid-treated membranes to capture gas phase NH_3(air)_, were deployed at sites GB and PC for periods of three and nine months (Figure S3). However, sufficient material was only available for isotopic analysis from 4 of the 12 samplers deployed (Table S2). In some cases, samplers became compromised by drip water that neutralized the filter and eluted trapped NH_4_+. In other cases, samplers remained intact but concentrations were low and isotope measurements were unreliable. The low concentrations of NH_4_^+^ that accumulated on the passive sampler membranes may reflect the effective scavenging by the acidic cave walls in the immediate environment. From the four passive samplers with sufficient material to measure N isotope ratios, those furthest from cave streams had δ^15^N_NHx_ values of -2.1‰ and -6.4‰, while samplers located just above streams had much lower δ^15^N_NHx_ values of -27.2‰ and -27.3‰ (Fig. 3a).

**Figure 3.**
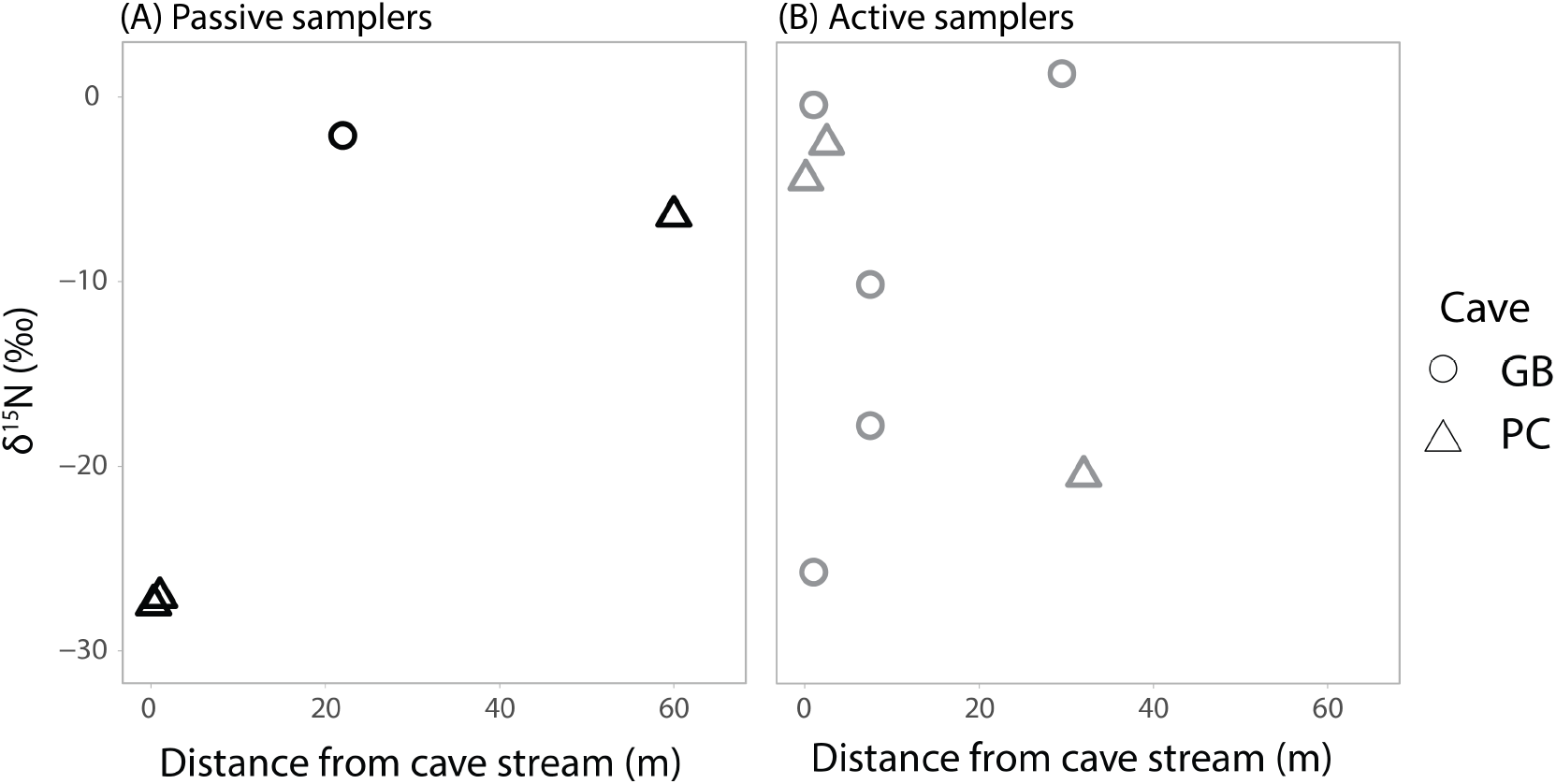
δ^15^N values of total dissolved nitrogen measured by passive (left) and active (right) ammonia samplers from sites GB and PC, versus distance from the nearest sulfidic stream.

Because most passive samplers did not recover enough NH_3_ for isotopic analysis, we also used active air samplers, which capture gas phase NH_3(g)_ and aerosol NH_4_^+^ as cave air is pumped through an acidified filter. Using active samplers, we recovered measurable NH_x_ from eight locations (Table S3). The δ^15^N values of actively sampled cave air varied considerably. The lowest δ^15^N value was -25.7‰ ± 4.3‰ from a sample collected just above the stream at site GB, although samples collected within 2.5 m of cave streams varied between -25.7‰ to -0.4 ‰ (Fig. 3). Samples collected 7.5 and 32 m from the stream had δ^15^N values as high as +1.3 ± 2.8‰ and as low as -20.5‰ ± 9‰ (Fig. 3).

We also measured the δ^15^N values of NH_4_^+^ in acidic condensation droplets collected from spiderwebs and snottites. These revealed clear spatial patterns, with very low δ^15^N_NH4_ values observed near streams, and higher values further away (Fig. 2). (Here, we use NH_4_^+^ rather than NH_x_ because, at very low pH, NH_3_ concentrations are vanishingly low [Eq. 5].) Droplet δ^15^N_NH4_ collected near the GB stream ranged from -29.0‰ to -24.6‰, while values in droplets 7.5 to 12.5 meters away were higher, ranging from δ^15^N_NH4_ of -12.6‰ to -4.7‰. The least acidic droplets (pH 4) collected 27.5 meters away yielded NH_4_^+^ with a still higher δ^15^N value of -3.8‰. At site RS, droplets collected at two locations 1.5 m from the stream had δ^15^N values of -22.6 and -2.5‰ (Fig. 2, Table S4).

### Other inorganic nitrogen sources

We evaluated other N sources by analyzing dissolved N compounds in drip pools and non-sulfidic streams fed by meteoric water rather than the sulfidic aquifer. Drip water pools had much lower NH_4_^+^ concentrations, with an average of 0.74 µM NH_4_^+^ in GB and 6.59 µM NH_4_^+^ in PC. We measured the isotopic composition of total reduced nitrogen (TRN) in these pools, δ^15^N_TRN_, which primarily reflects the ammonium but may also contain some fraction of dissolved organic nitrogen because analyses were conducted using persulfate oxidation. δ^15^N_TRN_ values were +0.3 to +4.7‰, except for one drip water pool at site PC that had a δ^15^N_TDN_ value of -30.9‰, likely impacted by droplets from nearby acidic walls (Table S5). Nitrate was not detected at GB or RS, although some drip pools at site PC contained 14-39 µM NO_3-_, which had δ^15^N_NO3_ values of -2.3 to +2.1‰. Samples collected from pools in higher (older) levels of the cave no longer influenced by hydrogen sulfide had nitrate concentrations as high as 227 µM NO_3_-, with δ^15^N_NO3_ values of +1.4 to +8.4 ‰. δ^18^O_NO3_ values from drip water pools in the lower levels ranged from -4.6 to -3.2‰, while drip water pools in the older upper levels of the cave had δ^18^O_NO3_ values ranging from +3.3 to -5.4‰ (Supplementary Table S5).

## Discussion

We found that ammonia gas in cave air and dissolved ammonium in acidic droplets are strongly depleted in ^15^N relative to ammonium in cave streams. The lowest δ^15^N_NH3_ values measured from passive and active filters near the streams were -27.3‰ and -25.7‰ (Figure 3), consistent with ammonium in nearby acidic droplets and organic N from wall biofilms that is similarly depleted in ^15^N (Fig. 1, 2). While a few samples did not follow this trend (discussed below), the presence of ^15^N-depleted ammonia and ammonium in the cave atmosphere and acidic biofilms is consistent with the hypothesis that volatilized ammonia is the primary or even sole nitrogen source for the acidic microbial communities and the invertebrates that feed on them (Fig. 4).

**Figure 4.**
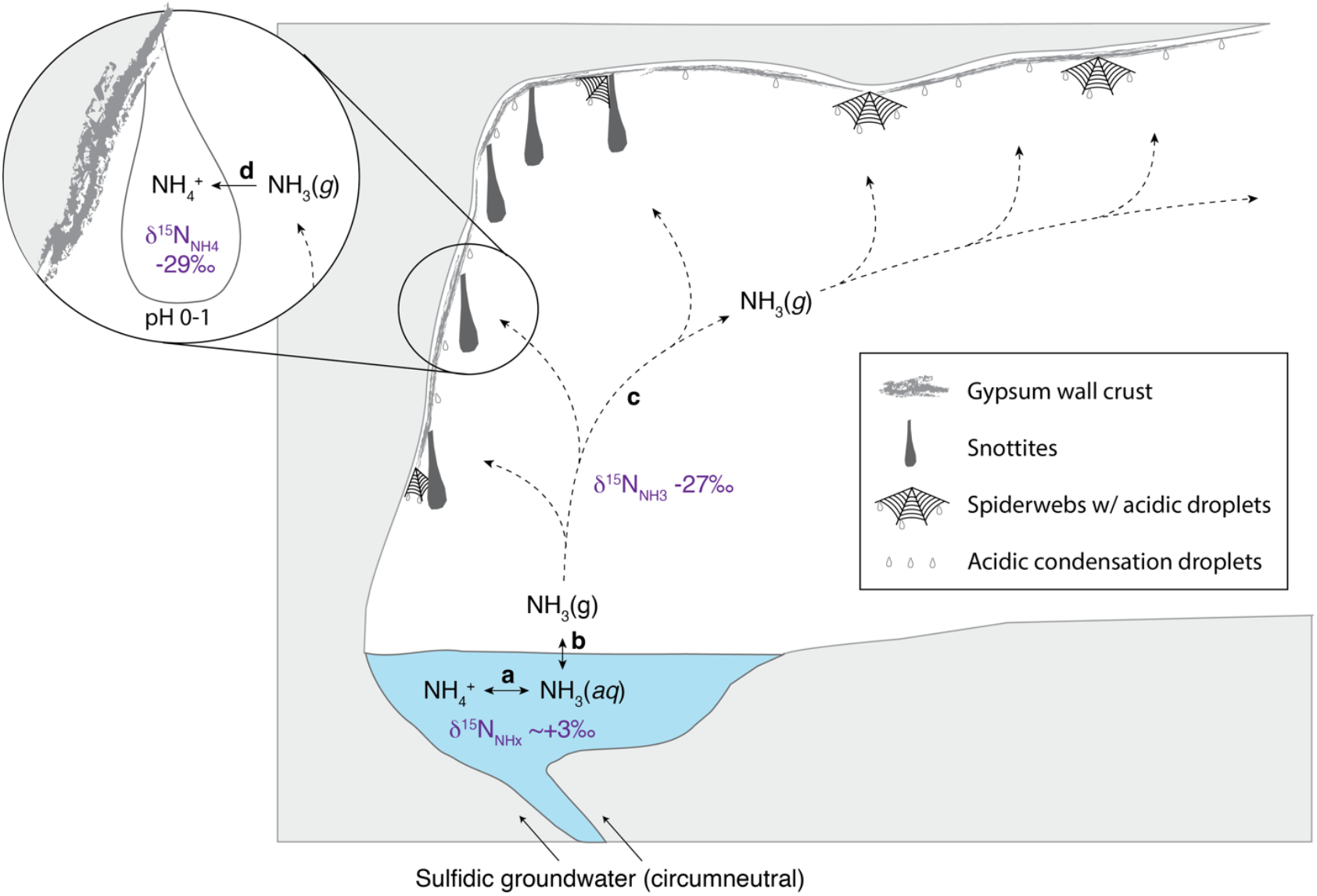
Summary of proposed ammonia volatilization and trapping processes resulting in extremely low δ^15^N values observed on cave walls. The lowest measured δ^15^N values are shown for NH_x_ in air and acidic droplets. Proposed reaction steps (bold letters) include **(a)** isotopic and chemical equilibrium between ammonium and ammonia in circumneutral streams (^15^ε ∼ 32‰; (42)), **(b)** isotopic and chemical equilibrium between dissolved ammonia and ammonia gas across the air-water interface (^15^ε ∼ 8‰; (43)), **(c)** possible fractionation during transport in cave air which may modulate expression of equilibrium isotope effects by counteracting equilibrium processes, and **(d)** near quantitative trapping in acidic snottites and condensation droplets on spiderwebs and gypsum surfaces (^15^ε ∼ 0‰).

### Isotope dynamics of ammonia volatilization and trapping in acidic droplets and biofilms

The presence of extremely low δ^15^N_NH4_ values in acidic droplets above the streams is consistent with NH_3(g)_ volatilization from the cave stream and reflects the large equilibrium isotope effect between NH_4_+_(aq)_ and NH_3(g)_, which results in δ^15^N_NH3_ values negatively offset from NH_4_^+^ at equilibrium by approximately 30‰ at 25°C (40–42). Superimposed on this effect is the equilibrium isotope effect between NH_3(aq)_ and NH_3(g)_, giving rise to an additional decrease of ∼8.3‰ in the δ^15^N of volatilized NH_3(g)_ relative to NH_3(aq)_ at equilibrium (40, 43). The combined effects thus yield NH_3(g)_ in the cave air that could be depleted in ^15^N by as much as 38‰ relative to the NH_4_^+^ pool in the stream.

The cave streams at Frasassi carry NH_x_ with an average δ^15^N of +3.3‰. It follows that dissolved NH_3_ (∼1% of the total NH_x_ pool at the pH of the stream waters) would exhibit δ^15^N values roughly 30‰ lower (∼ -26.7‰) and that volatilized NH_3(g)_ in the cave atmosphere would exhibit even lower δ^15^N values. Overall, this estimate is remarkably consistent with the lowest

δ^15^N values captured by the air sampler filters, and with condensation droplets in the high gas flux areas adjacent to the cave stream (δ^15^N as low as -29.1‰). This provides strong support for the hypothesis that volatilized NH_3(g)_ is the source of the very low δ^15^N values observed in organic matter above the water table (Fig. 1). Indeed, these patterns are also consistent with observations of ammonia volatilization in terrestrial environments from accumulated animal waste (e.g., confined feedlots, island bird colonies), where reported volatilization and transport of NH_3(g)_ can result in NH_3(g)_ that is strongly depleted in ^15^N relative to the source material (δ^15^N_NH3_ values 44‰ to 49‰ lower) (44) and in nearby plant matter being enriched in ^15^N (44– 49). In these cases, the isotopically light NH_3(g)_ is removed by volatilization, leaving a nitrogen pool with high δ^15^N values that are subsequently incorporated into plant biomass.

Capturing the light NH_3(g)_ signal on cave wall surfaces requires the NH_3(g)_ to dissolve and protonate into acidic droplets, effectively trapping the volatilized ammonia as NH_4_^+^. Consistent with this proposed mechanism, condensation droplets in the areas closest to degassing sulfidic cave streams with pH <1 contained 0.5 mM to 4.33 mM NH_4_^+^, despite cave atmosphere NH_3(*g*)_ concentrations ≤ 2.5 ppbv. The same condensation droplets exhibited very low δ^15^N values (Fig. 2, 3). In comparison, less acidic droplets further away from degassing streams contained only 18 µM NH_4_^+^, and freshwater drip pools had a maximum of 27.9 µM NH_4_^+^ (Fig. 2; Table S4).

### Possible explanations for higher δ^15^N_NHx_ values

NH_3(g)_ captured by passive samplers and some active samplers farther from the streams has relatively higher δ^15^N values (Fig. 3). There are guano deposits near cave entrances that have bulk δ^15^N values as high as +15‰ (Fig. 1; (27)). These guano deposits could supply NH_3(g)_ that is more enriched in ^15^N than the NH_3(g)_ volatilized from the sulfidic aquifer, and we attribute the higher δ^15^N values further from the streams (Fig. 3) to these guano deposits or other organic sources near cave entrances (Fig. 1). However, there was one sample collected 32 m from the water table at site PC with an δ^15^N_NHx_ value of -20.5‰, which indicates that depleted NH_3(g)_ from the streams can persist farther from the water table.

There were also two acidic droplet samples and several active air samples collected near streams that had NH_3(g)_ and NH4^+^ with relatively higher δ^15^N values (-10‰ to +1‰, Fig. 2, 3), and it is hard to explain these measurements by the ammonia volatilization mechanism described above. One possible explanation is that these higher δ^15^N values reflect NH_3(g)_ or NH_4_^+^ aerosols from other sources. In particular, NH_4_^+^ aerosols collected by active samplers could be more enriched in ^15^N compared to NH_3(g)_ if they originated from airborne, NH_4_^+^-bearing particulates or sources other than cave air NH_3(g)_. Another possibility is that the samples collected during extended pumping times could have been compromised by filter breakthrough, in which filter-bound NH_4_^+^ becomes enriched in ^15^N due to NH_3(*g*)_ loss through the filter as filter membranes become saturated. Indeed, the active sampler filters with higher NH4^+^ concentrations and, therefore, longer pumping times had higher δ^15^N values (Fig. S4). In all of these scenarios, the effect would be to enrich the trapped NH_4_^+^ in ^15^N relative to NH^3(*g*)^ in the cave air. Given these considerations, the lowest δ^15^N values we measured are more likely to represent the true isotopic composition of the NH_3(*g*)_ in the cave atmosphere.

### Microbial ammonia oxidation does not occur in the extremely acidic biofilms

Though it seems clear that the NH_4_^+^ found in acidic cave wall biofilms serves as a nutrient source, it is probably not used as an energy source by lithotrophic microorganisms despite very high NH_4_^+^ concentrations (Fig. 2a). Although ammonia-oxidizing bacteria (AOB) and archaea (AOA) are widely distributed in many natural environments, including marine (50), soil (51–53), freshwater (54), and extreme environments such as low pH hot springs (55–58), there is no evidence for microbial ammonia oxidation in the acidic areas of the cave system. While Gibbs energy estimates indicate that ammonia oxidation to nitrite (NO_2_^-^) or nitrate (NO_3_^-^) is energetically favorable in the low pH snottites and condensation droplets (Fig. S5), previous investigations using both metagenomics and rRNA gene surveys have yielded no evidence for AOB or AOA in extremely acidic snottite biofilms (19, 20, 30, 39) or on nearby acidic gypsum surfaces (59, 60). One potential explanation may be the presence of sulfide and other sulfur compounds, which have been shown inhibit ammonia monooxygenase (AmoA) (61–66). However, archaeal *amoA* sequences have been detected in some sulfidic environments, such as estuarine sediments (67) and sulfidic circumneutral cave stream biofilms in Movile Cave (68). Instead, ammonia oxidation may be inhibited by the extremely low abundance of NH_3_ (presumed to be the active substrate) at the relevant pH values (Eq. 5; pKa = 9.25). While ammonia oxidation has been demonstrated in culture from pH 4 (52) to 2.5 (69) and in the environment at pH values as low as 2.4 (55, 70), to our knowledge no activity has been documented at pH values that typify the Frasassi wall biofilms (pH < 2).

## Conclusions

Our isotopic and ammonium concentration data support the hypothesis that, despite very low NH_3(g)_ concentrations in cave air, lithoautotrophic communities on extremely acidic cave walls rely mainly or exclusively on volatilized NH_3(g)_ as an N source. The nitrogen isotopic values we measured are consistent with prior experimental results (40, 43) and observations from other terrestrial environments (44–49). Our results are also consistent with extremely low δ^15^N values found in biofilms adjacent to Frasassi cave streams in previous work (19, 27, 28, 38) (Fig. 1b), as well as with the absence of any evidence for nitrogen fixation or ammonia oxidation in Frasassi snottites (30, 31). Although NH_3(g)_ with very low δ^15^N values persists in some areas distal to sulfidic streams, most cave wall surfaces in these areas are not acidic enough to serve as effective ammonium traps. Higher δ^15^N_NH3_ values and lower NH_x_ concentrations in cave air and water farther away from sulfidic streams suggest a greater influence of other nitrogen sources in areas of the cave that are not directly in the path of the gas fluxes from the streams.

This work provides the first direct evidence for subaerial lithoautotrophic ecosystems that rely on volatilized ammonia as a nitrogen source. This unique situation arises from the synergy between the metabolic activity of sulfide oxidizing microbial biofilms that produce extremely acidic conditions, which in turn promote effective trapping of volatilized ammonia. Further from cave streams, cave wall surfaces and biofilms are buffered by limestone host rock and become circumneutral, decreasing the efficacy of this natural acid trap and requiring increased dependence on other N sources or slower growth. Our work highlights the ways in which atmospheric delivery of nutrients and energy resources can support life, and the strong controls airborne compounds exert on the cave ecosystem. Sulfidic caves showcase a novel mechanism for nutrient acquisition and subsurface lithotrophic primary production that may enhance the habitability of natural and engineered systems on Earth and other planetary bodies.

## Materials and Methods

### Sample Collection

Samples were collected during October 2021 and July 2022 at three passages within the larger Frasassi cave system where turbulent sulfidic streams are accessible via technical caving routes: Grotta Bella (GB), Pozzo dei Cristalli (PC), and Ramo Sulfureo (RS) (Fig. S2). At each site, both passive and active techniques were used to sample cave air NH_3_ at varying distances from the stream, to evaluate spatial patterns in concentration and isotopic composition with proximity to the source waters. At the time of collection, concentrations of O_2_, H_2_S, CO_2_, SO_2_, CO, and CH_4_ in the cave air were measured with ENMET SAPPHIRE and ENMET RECON/4a portable gas detectors (ENMET, Ann Arbor, MI, USA). pH of drips was measured using pH paper (ranges 0-2.5, 4-7, or 2-9).

Passive gas samplers (ALPHA Samplers, UK Centre for Ecology & Hydrology, Bangor, UK) were loaded with citric acid-treated filters (25mm) situated inside small plastic housings with permeable covers designed to acid-trap gas phase NH_3_ (71).Triplicate housings were attached to a small plastic mount and suspended using monofilament. After passive sampler deployments of 3 to 9 months, filters were rinsed with DI water to elute all trapped NH_4_^+^. Low concentrations required pooling of all three passive gas sampler filters from each site.

Active aerosol samplers were designed following previously described sampling protocols (42) and employed a similar acid-trapping approach as used for the passive gas samplers. A small 12V battery powered vacuum pump (KNF Neuberger, Inc., Trenton, NJ, USA) was used to pull cave air through a citric acid-coated filter mounted in an acrylic filter housing (47mm) at a rate of 7.5-9 L/min as monitored by flow meter. Samplers were generally operated for 0.5-2 hours per deployment. Once completed, filters were removed from the housing and placed into 10ml MilliQ water to elute all NH_4_^+^. Sampling periods and flow rates are presented in the supplementary materials (Table S3). Concentrations of the eluted NH_4_^+^ were measured as described below and used together with the total volume of air sampled to calculate concentrations in the cave air.

Dissolved oxygen, pH, oxidation-reduction potential, and specific conductivity of cave streams and drip pools were determined with ProSolo and ProPlus multimeters (YSI Incorporated, Yellow Springs, OH, USA), or, for shallow or small volume dripwaters, compact pH and conductivity meters (LAQUAtwin pH-33 and EC-11, Horiba, Kyoto, Japan). Water samples for dissolved sulfide analysis were filtered using 0.2 μM polyethersulfone membranes and preserved in zinc chloride (ZnCl_2_) for sulfide analysis with the methylene blue method using Hach sulfide reagents 1 and 2 (Hach method 8131 [Hach Company, Loveland, CO, USA], equivalent to US EPA Method 376.2).

### Ammonium concentrations

Ammonium concentrations were measured by flow injection analysis using a diffusion cell and conductimetry (72). Sample water (0.5ml) was manually injected and introduced into a stream of 10 mM NaOH and 0.2M Na-citrate flowing opposite a stream of 50 μM HCl separated by a strip of Teflon housed in a 25 cm diffusion cell. On contact with the base, all ammonium is rapidly converted to ammonia, which diffuses from the NaOH to the HCl side causing a quantifiable change in conductivity. Ammonium standards were regularly measured for establishing a calibration curve, and all samples were measured in duplicate or triplicate. Samples with high NH_4_^+^ were diluted in DI water as needed.

### Nitrate concentrations

Nitrate concentrations of stream waters and dripwater pools were measured using the denitrifier method (described below), wherein nitrate is quantitatively converted to N_2_O and quantified on an Isoprime 100 isotope ratio mass spectrometer (IRMS). Integrated N_2_O peak areas (m/z 44) of samples were compared to injections of known nitrate reference materials and divided by injected volumes for calculation of sample nitrate concentration.

### Isotopic composition

All nitrogen species in aqueous samples (eluted passive filters, eluted aerosol filters, directly collected water samples, as well as spiderweb and snottite droplets) were first converted to N_2_O for N (and O, if applicable) isotope analysis using an Isoprime100 isotope ratio mass spectrometer (IRMS) in the Wankel Lab at WHOI. Samples for NO_3_^-^ isotope analysis were converted using the denitrifier method, wherein 20 nmoles of NO_3_^-^ is quantitatively converted to N_2_O by a culture of denitrifying bacteria in 20ml headspace vials (73, 74). The evolved N_2_O is then quantitatively purified using a customized purge and trap system before being introduced into the IRMS. Nitrate δ^15^N and δ^18^O were corrected for instrument drift and linearity by regular analyses of isotope reference materials USGS 32, USGS 34 and USGS 35 (75).

Analyses of δ^15^N of NH_4_^+^, including those from air samplers, cave waters and droplets collected from the cave walls, were first converted to NO_3_^-^ by persulfate oxidation before being converted to N_2_O using the denitrifier method as described above (76). Reference NH_4_^+^ standards USGS 25 and USGS 26 were used to correct for contribution of any blanks arising from the potassium persulfate reagent. While this method captures the δ^15^N of the total N pool (DIN + DON), for all samples of cave air and wall droplets total N was comprised of >99% NH_x_ and thus reported values accurately reflect the δ^15^N of NH_x_. In cave streams and ponded waters, where NO_3_^-^ was also present in the sample, the δ^15^N of the total reduced N pool (e.g., TRN = NH_4_^+^ + DON) was calculated by isotope mass balance.

### Sources of error

The filters used in both the passive and the active samplers were coated with citric acid following procedures outlined in (77), who reported a δ^15^N precision of ±1.6‰ (2σ) and an operative capacity of ∼400µg NH_3_ at concentrations ≤ 207 ppb_v_. Using the same filters to collect particulate NH_4_^+^ (42) reported a δ^15^N precision of ±0.9‰ (1σ). All NH_3_ and NH_4_^+^ collected on these filters was eluted into MQ water by sonication for 60 minutes, filtered (0.2µm) and then frozen prior to analysis. Our filter samples were treated in the same way with the exception of sonification. The precision of δ^15^N measurements using the denitrifer method (73) based on replicate analysis of standard reference materials is ±0.3‰ (1σ).

### Henry’s law and Gibbs energy calculations

Expected NH_3_(g) concentrations in cave air at equilibrium with stream waters were calculated using PHREEQC v3 in the package “phreeqc” v. 3.7.5 (78) in R (79). Calculations used the PHREEQC database as well as values for pH, temperature, and NH_4_^+^(*aq*) concentration values measured in this study, and major ion concentrations for the PC, RS, and GB streams reported in (21). NH_3_(g) equilibrium values in Table S1 are reported as concentrations in ppbv by assuming a fugacity constant of 1. Gibbs energies for the complete oxidation of ammonia to nitrate (performed by COMAMMOX microorganisms) as well as oxidation of ammonia to nitrite (catalyzed by AOA and AOB) were performed in CHNOSZ (80) as in (81). Gibbs energies were calculated as a function of pH and ammonium concentration, assuming a temperature of 25°C, 20.9% oxygen, and activities of nitrate and nitrite of 10^-12^.

## Acknowledgments

We thank A. Montanari, P. Metallo, and L’Associazione Le Montagne di San Francesco for logistical support and the use of facilities and laboratory space at the Osservatorio Geologico di Coldigioco. Thanks to H. Aronson for assistance with Gibbs energy calculations in CHNOSZ, and to Z. Havlena, K. Green, E. Barlow, N. Bice, and A. Wankel for field assistance and sample collection.

This research was primarily supported by the NASA Exobiology program (award 80NSSC20K0619) to D.S.J., S.D.W., H.VG. and J.C.S., with additional support from the New Mexico Space Grant Consortium to M.B.B.

Sandia National Laboratories is a multi-mission laboratory managed and operated by National Technology & Engineering Solutions of Sandia, LLC (NTESS), a wholly owned subsidiary of Honeywell International Inc., for the U.S. Department of Energy’s National Nuclear Security Administration (DOE/NNSA) under contract DE-NA0003525. This written work is authored by an employee of NTESS. The employee, not NTESS, owns the right, title, and interest in and to the written work and is responsible for its contents. Any subjective views or opinions that might be expressed in the written work do not necessarily represent the views of the U.S. Government. The publisher acknowledges that the U.S. Government retains a non-exclusive, paid-up, irrevocable, world-wide license to publish or reproduce the published form of this written work or allow others to do so, for U.S. Government purposes. The DOE will provide public access to results of federally sponsored research in accordance with the DOE Public Access Plan.

## Notes

### Competing Interest Statement

The authors have declared no competing interest.

