## Supplementary Figures for "Volatilized ammonia supports extremophilic cave ecosystems with unusual nitrogen isotopic signatures"

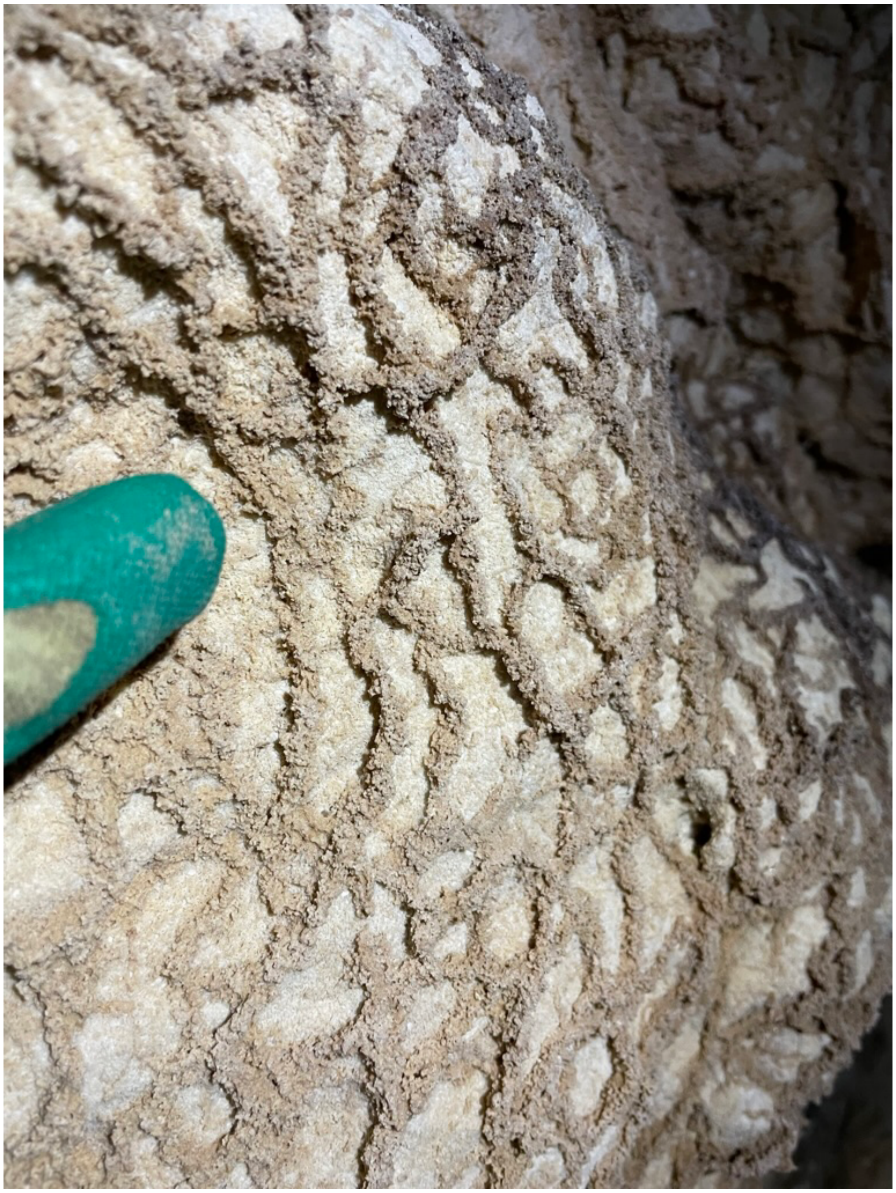

**Supplementary Figure S1**. Biovermiculations in Grotta Bella.

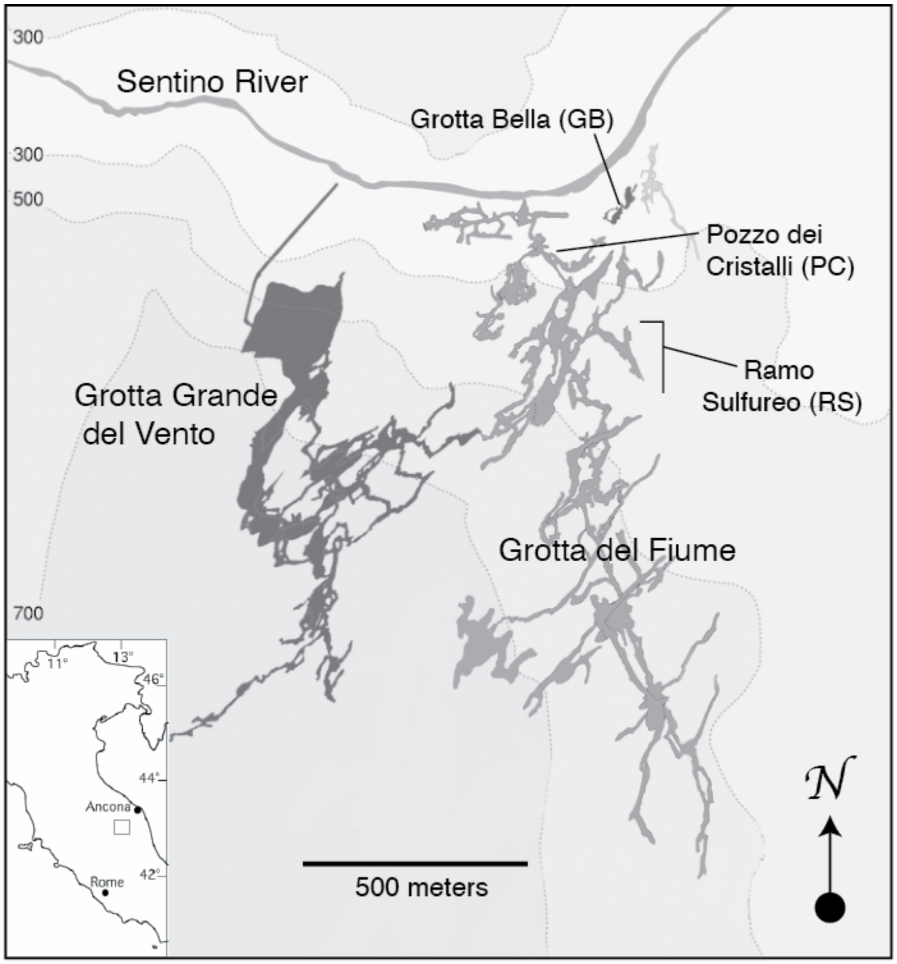


**Supplementary Figure S2**. Map of the Frasassi cave system, showing the three sampling locations used in this study. Base map courtesy of the Gruppo Speleologico CAI di Fabriano.

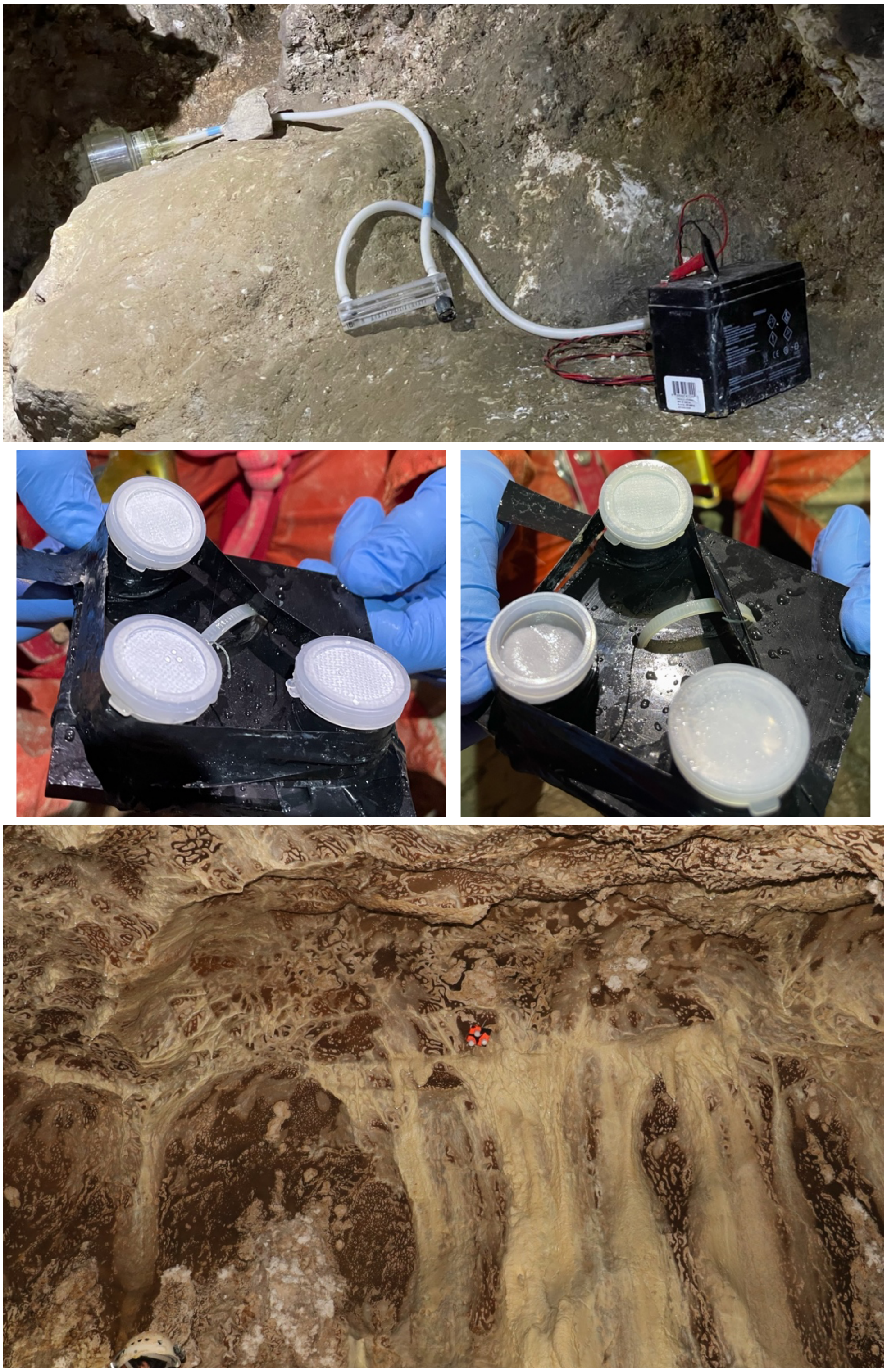

**Supplementary Figure S3**. Active (top) and passive (bottom three images) cave air samplers. In the passive samplers, the acid-treated filter is protected by a gas-permeable cover; image in the middle right shows one that was wet upon collection and the bound ammonium likely lost. The bottom image shows a deployed passive ammonia sampler.

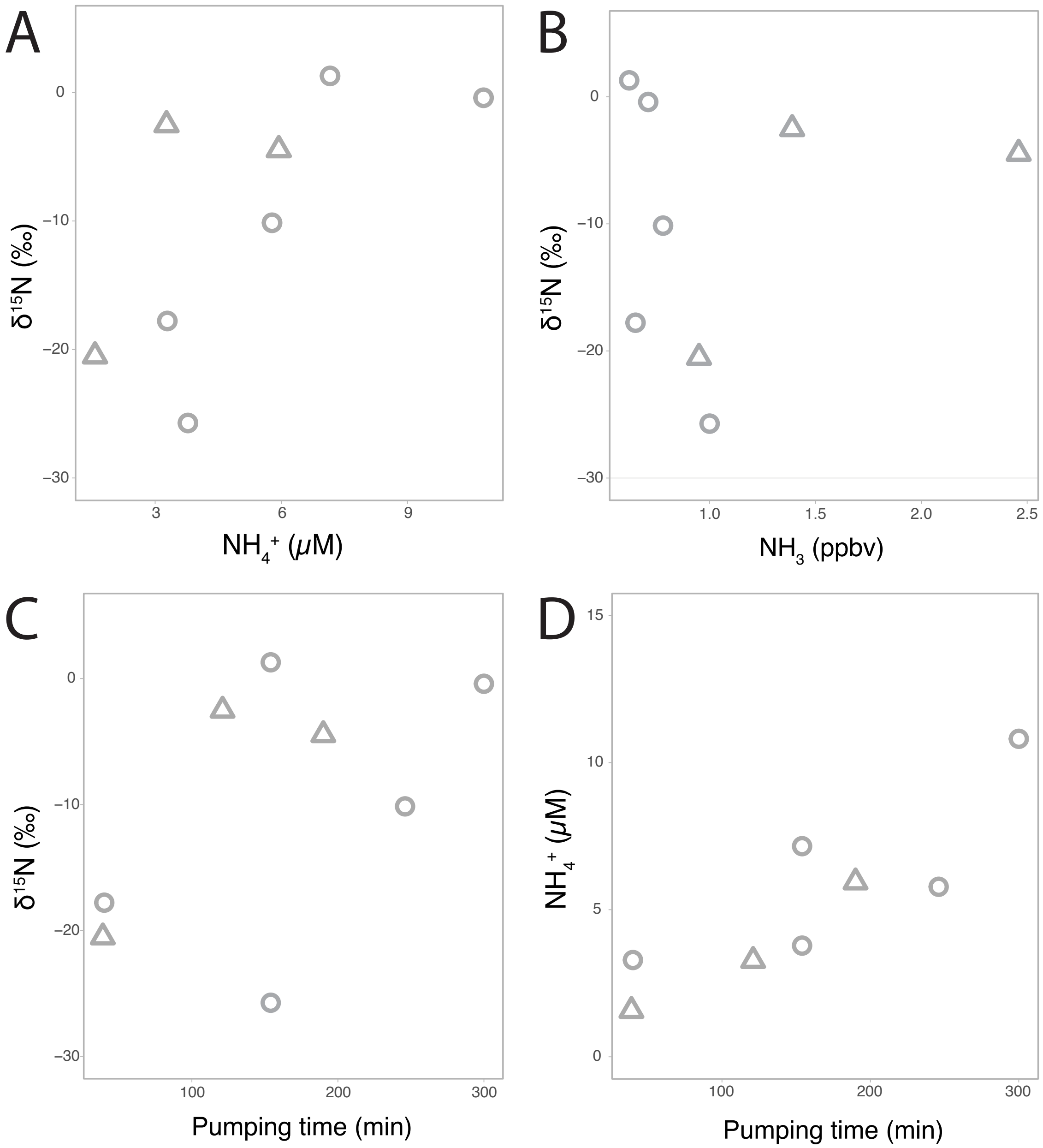


**Supplementary Figure S4**. $\delta^{15}N$ of ammonium recovered from active cave air samplers versus (A) the concentration of ammonium concentration eluted from filters, (B) the calculated concentration of $NH_3$(g) in the cave air, and (C) pumping time. Panel D shows the concentration of ammonium concentration eluted from filters versus pumping time. Circles indicate samples from site GB, and triangles indicate samples from site PC.

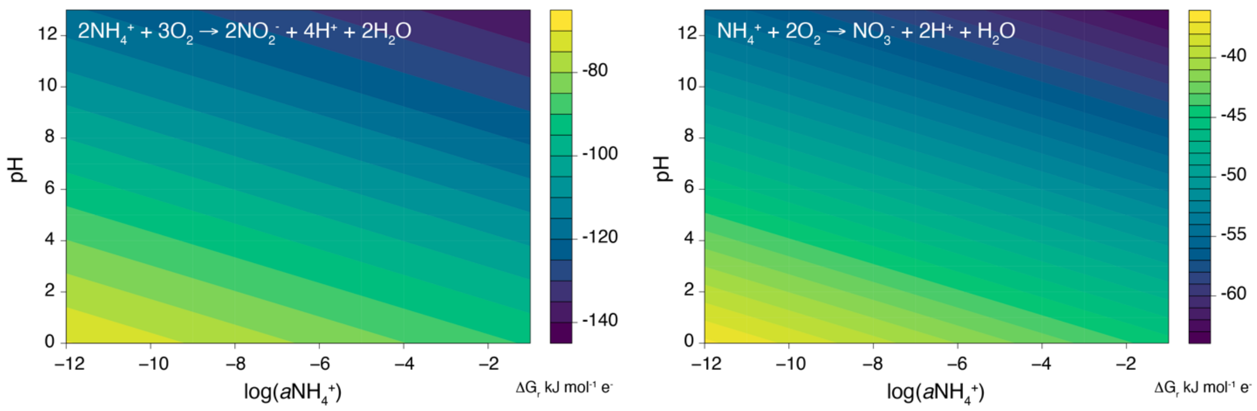


**Supplementary Figure S5.** Gibbs free energy of ammonium oxidation to nitrite and nitrate as a function of pH and ammonium concentration, assuming a temperature of 25°C, 20.9% oxygen, and activities of nitrate and nitrite of $10^{-12}$.
